# Loss and gain of function experiments implicate TMEM18 as a mediator of the strong association between genetic variants at human Chromosome 2p25.3 and obesity

**DOI:** 10.1101/122853

**Authors:** Rachel Larder, M. F. Michelle Sim, Pawan Gulati, Robin Antrobus, Y.C. Loraine Tung, Debra Rimmington, Eduard Ayuso, Joseph Polex-Wolf, Brian Y.H. Lam, Cristina Dias, Darren W. Logan, Sam Virtue, Fatima Bosch, Giles S.H. Yeo, Vladimir Saudek, Stephen O’Rahilly, Anthony P. Coll

**Affiliations:** University of Cambridge Metabolic Research Laboratories, Level 4, Wellcome Trust-MRC Institute of Metabolic Science, Box 289, Addenbrooke’s Hospital, Cambridge, CB2 0QQ. United Kingdom; Cambridge Institute for Medical Research, University of Cambridge, United Kingdom; Center of Animal Biotechnology and Gene Therapy and Department of Biochemistry and Molecular Biology, School of Veterinary Medicine, Universitat Autònoma de Barcelona, 08193-Bellaterra and Centro de Investigación Biomédica en Red de Diabetes y Enfermedades Metabólicas Asociadas (CIBERDEM), Spain; Wellcome Trust Sanger Institute, Wellcome Genome Campus, Hinxton-Cambridge, CB10 1SA, United Kingdom.

## Abstract

An intergenic region of human Chromosome 2 (2p25.3) harbours genetic variants which are among those most strongly and reproducibly associated with obesity. The molecular mechanisms mediating these effects remain entirely unknown. The gene closest to these variants is *TMEM18*, encoding a transmembrane protein localised to the nuclear membrane. The expression of *Tmem18* within the murine hypothalamic paraventricular nucleus was altered by changes in nutritional state, with no significant change seen in three other closest genes. Germline loss of *Tmem18* in mice resulted in increased body weight, which was exacerbated by high fat diet and driven by increased food intake. Selective overexpression of *Tmem18* in the PVN of wild-type mice reduced food intake and also increased energy expenditure. We confirmed the nuclear membrane localisation of TMEM18 but provide new evidence that it is has four, not three, transmembrane domains and that it physically interacts with key components of the nuclear pore complex. Our data support the hypothesis that *TMEM18* itself, acting within the central nervous system, is a plausible mediator of the impact of adjacent genetic variation on human adiposity.

## INTRODUCTION

The understanding of human obesity has benefited greatly from advances in molecular genetics. In addition to the identification of many mechanistically illuminating, highly penetrant monogenic disorders, genome-wide association studies (GWAS) have identified multiple common genetic variants strongly associated with body mass index (BMI) (1-5). Many of these loci are within, or close to, genes which to date, have not been recognised to encode proteins with a role in the control of energy homeostasis. Of note, these genes show a strong preponderance to being highly expressed in the central nervous system, a finding which is congruent with the fact that monogenic disorders leading to obesity largely exert their effects through a disruption of the central control of appetite and energy balance (6).

To fulfil the potential of these findings and to gain a better understanding of the underlying circuitry involved in the control of human energy balance, there is an imperative to bridge the gap between the identification of a variant as being associated with an adiposity phenotype and the understanding of how that variant actually influences energy balance. Murine models have proven to be essential in moving from statistical association to a fuller understanding of the biology of the genes potentially regulated by these genetic variants. For example, the robust correlation between BMI and polymorphisms in the human fat mass and obesity-associated (*FTO*) gene has since been followed by a number of rodent studies which have demonstrated a role for both *FTO* and its neighbouring genes in energy homeostasis and body composition (7-13).

A strong association between increased BMI and a region of human chromosome 2, near to the gene *TMEM18*, has been repeatedly demonstrated in both adults and children (2, 14-18). *TMEM18* encodes a small (140 amino acid) protein reported to be comprised of three transmembrane domains (19). Like *FTO, TMEM18* had not been recognised as having a role in energy homeostasis prior to its identification by GWAS and relatively little is known about its function, save that it is expressed in the brain (19) and may act as a DNA-binding protein (20). The protein is highly conserved across several species (19), Wiemerslage *et. al*. recently reported that loss of TMEM18 from insulin producing cells in *Drosophila* induces a metabolic state resembling type 2 diabetes (21).

We have therefore undertaken a range of studies to determine whether TMEM18 plays a role in the control of energy balance in mammals.

## RESULTS

### *Tmem18* expression within the hypothalamic paraventricular nucleus is nutritionally regulated

We initially examined the expression pattern of *Tmem18* in a range of murine tissues and two mouse hypothalamic cell lines (N-46 and GT1-7). In keeping with previous reports (19), *Tmem18* was widely expressed, with expression observed in several regions of the brain, in brown adipose tissue and in both hypothalamic cell lines (Supplementary Fig 1). To define the expression pattern within the hypothalamus, we used immunohistochemistry and found TMEM18 to be enriched within the paraventricular nucleus (PVN) and arcuate nucleus (ARC) (Supplementary Fig 1). Following this, we re-interrogated an existing data set arising from detailed transcriptomic analysis of distinct hypothalami nuclei we had previously acquired by laser capture microdissection (LCM) (22). This was based on hypothalamic tissue from 3 groups of mice (*ad lib* fed, 48 hr fasted and fasted for 48 hrs with leptin administered throughout the fast). Within the ARC, microarray analysis showed no change in *Tmem18* expression level across the three different nutritional states (data not shown). We therefore focused further on the PVN and repeated the fed, fasting, fasting plus leptin experiment in a second, separate cohort of mice, undertaking qPCR on laser capture PVN tissue. 48 hr fasting decreased PVN *Tmem18* expression by 70% with leptin administration during the fast restoring expression back to fed levels (Fig 1).

**Figure 1.**
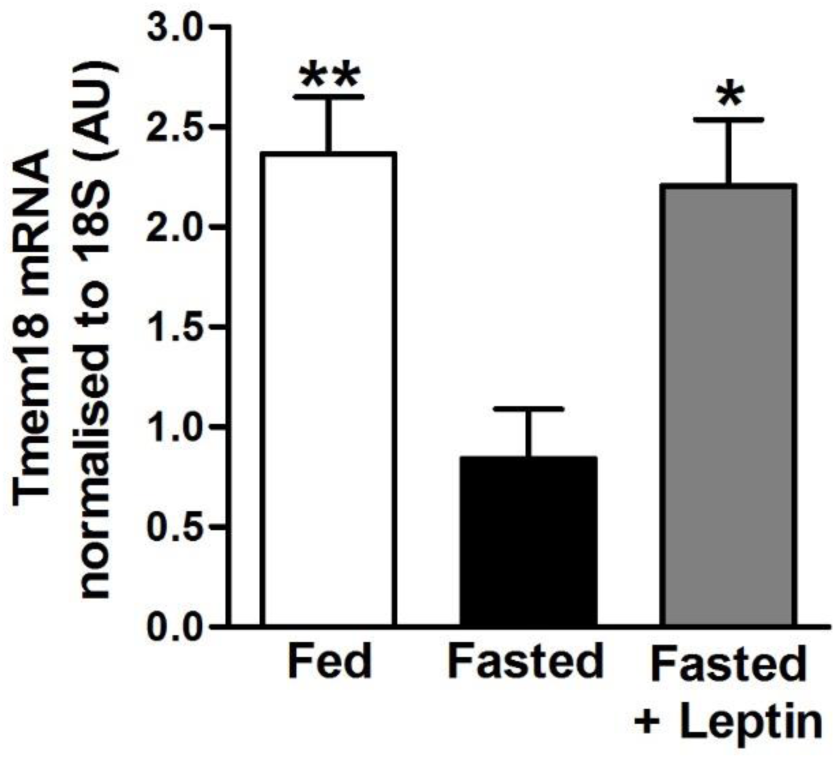
*Tmem18* expression within the hypothalamic paraventricular nucleus is nutritionally regulated. Q-RT-PCR analysis showing changes in *Tmem18* gene expression in the PVN of wildtype mice that have been fed, fasted for 48hrs or fasted for 48hrs with leptin administration.

In a further set of animals, we undertook transcriptomic analysis (RNA Seq) of 4 distinct hypothalamic nuclei (arcuate, ventral medial, paraventricular and dorsal medial nuclei) in tissue acquired by LCM from wildtype mice who had *been ad libitum* fed or fasted for 24hrs (GEO accession number GSE96627). Analysis of control genes, in each of the nuclei, revealed expected results indicating that good microdissection had been achieved and the animals had responded appropriately to the fast (Supplementary Fig 2). *Tmem18* expression was observed in all the four nuclei examined but differences in fed *vs* fast were only seen in the PVN (increased by ∼30%) (Supplementary Fig 3). Of note, there was no significant change in fed *vs* fasted expression of vicinal genes (*Sh3yl1, Snth2* and *Acp1*) in any of the 4 nuclei studied (Supplementary Fig 4). Thus, *Tmem18* expression in the PVN appears to be altered during feeding and fasting in a biphasic manner with an initial modest rise followed by a more profound fall, which is dependent on a reduction in circulating leptin.

### Loss of Tmem18 expression results in an increase in body mass

Knockout mice carrying mutant allele *Tmem18^tm1a^*^(^*^EUCOMM^*^)^*^Wtsi^* (abbreviated to *“Tmem18^tm1a^”*) were generated on a C57BL/6 genetic background as part of the European Conditional Mouse Mutagenesis Program (EUCOMM) (23). The introduction of the allele results in targeted disruption of exon 2 of *Tmem18*. Generation of hypomorphic mice, as well as the disruption of neighboring genes, has previously been demonstrated using the “knockout first” strategy (24-27). We therefore determined the expression levels of *Tmem18*, and the genes that surround it on mouse chromosome 12, by Q-RT-PCR analysis in *Tmem18^tm1a^* mice and their wildtype and heterozygous littermates. Data revealed a very low level of residual *Tmem18* transcript within the hypothalamus of homozygous mice (2.1% ±1.4, Supplementary Figure 5A), whilst heterozygous mice demonstrated a 50% decrease in transcript expression compared to *Tmem18^+/+^* (49% ±9, Supplementary Figure 5A). *Tmem18* was the only transcript at that locus to be altered by introduction of the allele (Supplementary Figure 5B) suggesting that there were no local ‘off target’ effects. The mice were viable with expected homozygous mutant offspring born from heterozygous crosses. WT and homozygous *Tmem18^tm1a^* mice were weaned onto a diet of normal chow and weighed weekly until 16 weeks of age. Female *Tmem18^tm1a^* homozygotes showed no differences in body weight or body composition compared to wildtype littermates (Supplementary Figs 6A, C and E). Male *Tmem18^tm1a^* homozygotes, on normal chow, had significantly increased body weight by 14 weeks of age (Fig 2A). This increase in body weight was due to a significant increase in both fat and lean mass (Fig 2C). At 16 weeks of age male homozygotes weighed on average 1.9 g more than wildtype littermates and had both increased gonadal white adipose tissue (gWAT) and brown adipose tissue (BAT) (Fig 2E).

**Figure 2.**
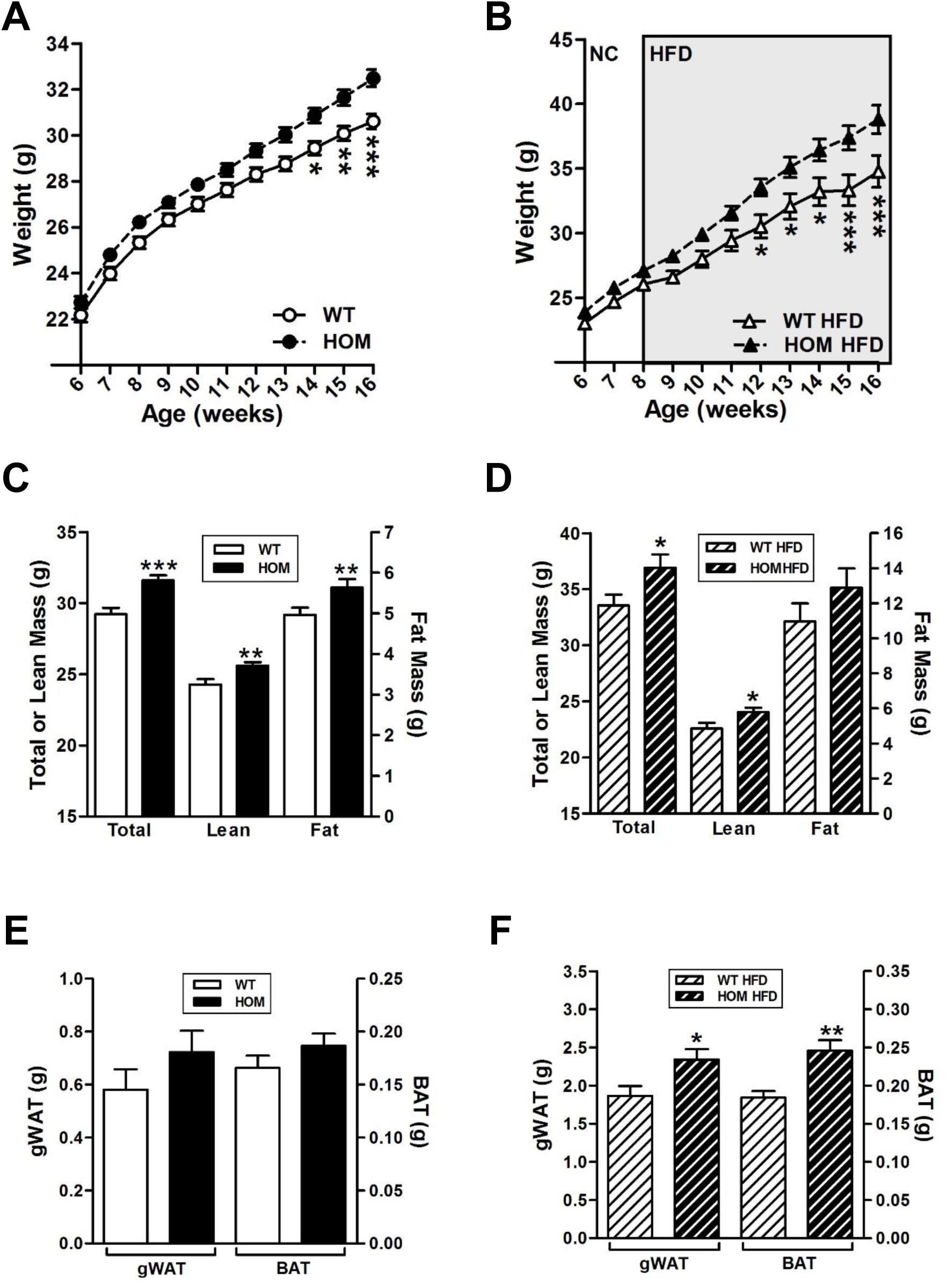
Effect of loss of expression of <u>Tmem18</u> on body weight and body composition in male mice maintained either on a normal chow diet or 45% HFD. **(A)** Body weights of *Tmem18^wt/wt^* (WT) and *Tmem18^tm1a/tm1a^* (HOM) male mice on normal chow (WT, n=23; HOM, n=23). **(B)** Body weights of WT and HOM male mice placed on a high fat diet (HFD) at 8 weeks of age (WT, n=15; HOM, n=20). **(C)** Body composition analyses showing total, lean or fat mass of 14-week old WT and HOM male mice on normal chow (WT, n=16; HOM, n=16). **(D)** Body composition analyses showing total, lean or fat mass of 14-week old WT and HOM male mice placed on HFD at 8 weeks of age (WT, n=18; HOM, n=20). **(E)** Weights of gonadal white adipose tissue (gWAT) and brown adipose tissue (BAT) of WT and homozygous mice at 18 weeks of age (WT, n=12; HOM, n=9). **(F)** Weights of gonadal white adipose tissue (gWAT) and brown adipose tissue (BAT) of 18-22 week old WT and homozygous mice fed a HFD from 8 weeks of age (WT, n=15; HOM, n=20). Data are expressed as mean ± SEM. WT mice are represented by white symbols and bars. HOM mice are represented by black symbols and bars. Time-course data were analysed using the repeated measures ANOVA model. p-values for C-F were calculated using a two-tailed distribution unpaired Student’s t-test and established statistical significance as follows * p<0.05, ** p<0.01, *** p<0.001 vs WT mice.

### Male *Tmem18* deficient mice fed a high fat diet become more obese because of an increase in food intake

To determine the effects of a high fat diet on weight gain in *Tmem18* deficient mice, WT and homozygous *Tmem18^tm1a^* mice were weaned onto a normal chow diet at 3 weeks of age then switched to a 45% fat diet (HFD) at 8 weeks of age. All mice were weighed weekly from weaning until 16 weeks of age. Female *Tmem18^tm1a^* homozygotes on a HFD showed no differences in body weight or body composition compared to wildtype littermates (Supplementary Fig 6B, D and F). However, male *Tmem18^tm1a^* homozygotes on a HFD displayed significantly increased body weight by 12 weeks of age (Fig 2B). This increase in body weight was a result of an increase in both fat and lean mass (Fig 2D). At 16 weeks of age male homozygotes on a HFD weighed ∼4.1 g heavier than wildtype littermates and had both increased gWAT and BAT (Fig 2F). Food intake in these mice was assessed on two occasions, firstly over a 10-day period at 15 weeks of age whilst individually housed in home cages (Fig 3A), and secondly over the 72hr period at 16 weeks of ages when mice were being analyzed for energy expenditure (Fig 3B). On both occasions, *Tmem18* deficient mice consumed significantly more energy than wildtype littermates. Interestingly, at the time of indirect calorimetry, *Tmem18* deficient males fed a high fat diet displayed a significantly increased energy expenditure when compared to wildtype littermates (ANCOVA analysis with correction for body weight differences, *p=0.015, Figs 3C and 3D). Of note, Q-PCR analysis detected no significant difference in the expression of various markers of thermogenesis in brown adipose tissue between the two genotypes (Fig 3E) and analysis of beam break data for the period when the mice were in the indirect calorimetry machine (72hrs) revealed no difference in activity levels between the two genotypes (Fig 3F). Thus, the increase in food intake (13.9%) in animals lacking *Tmem18* was partially compensated for by a rise in energy expenditure (7.0%) which explains the modest increase in weight.

**Figure 3.**
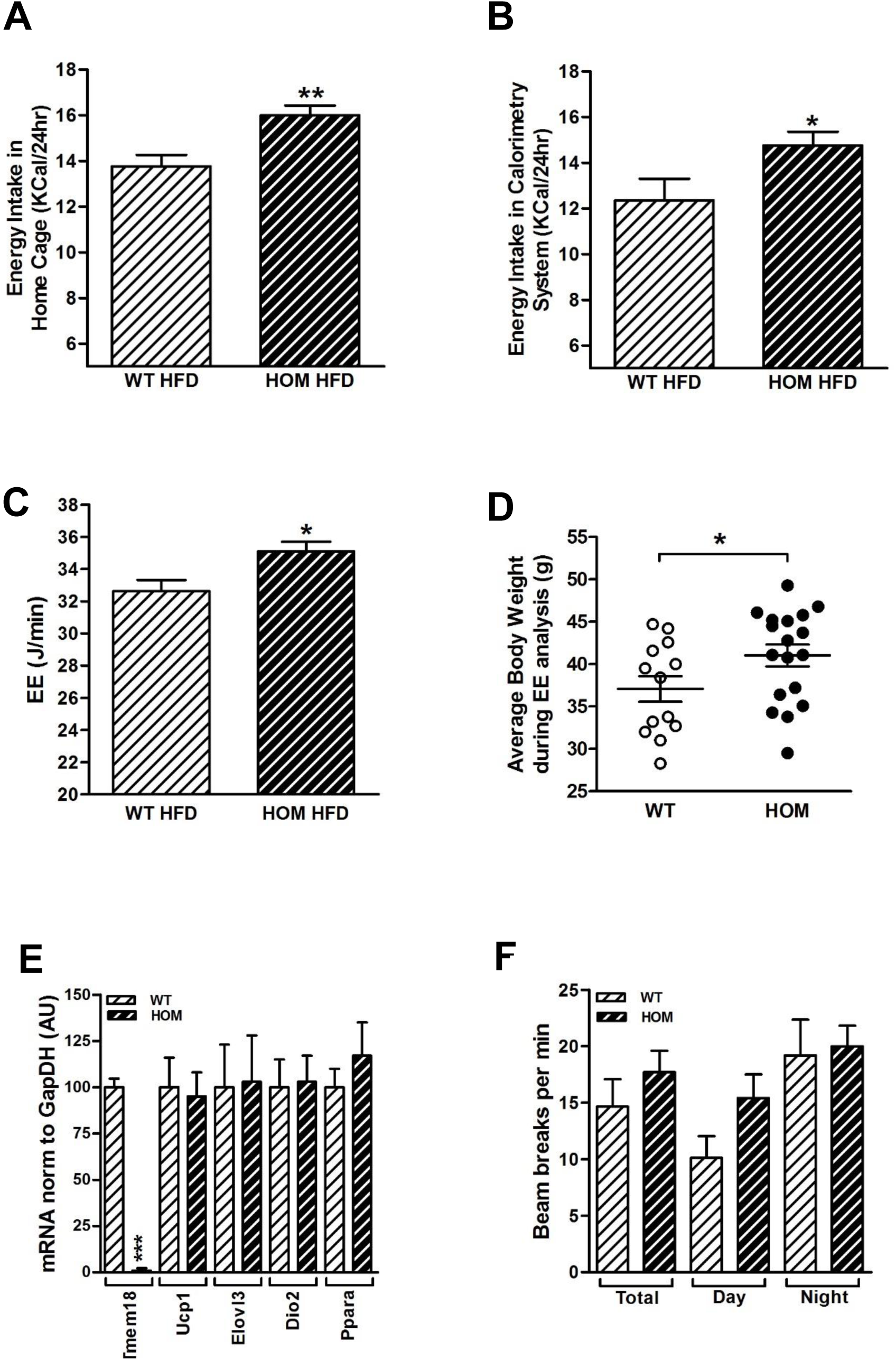
Effect of loss of expression of *Tmem18* on energy homeostasis, activity levels and food intake in male mice fed a HFD. **(A)** Average 24-hour energy intake of WT and homozygous male mice, measured in their home cage (WT, n=13; HOM, n=18). **(B)** Average 24-hour food intake of WT and homozygous male mice, measured in the indirect calorimetry cage (WT, n=13; HOM, n=18). **(C)** ANCOVA analysis of 24-hour energy expenditure (EE) with body weight at time of analysis (WT, n=13; HOM, n=18). **(D)** Average bodyweight of individual animals during 72hr analysis of energy expenditure (WT, n=13; HOM, n=18). **(E)** Q-RT-PCR analysis of *Tmem18* and brown adipose tissue (BAT) markers *Ucp1, Elovl3* and *Ppara* expression in WT and HOM male mice (WT, n=7; HOM, n=5). **(F)** Activity of WT and HOM mice during indirect calorimetry experiment represented by beam breaks per minute during total time in cage, the day and at night (WT, n=13; HOM, n=18). Data are expressed as mean ± SEM. p-values were calculated using a two-tailed distribution unpaired Student’s t-test and linear regression and established statistical significance as follows * p<0.05, ** p<0.01, *** p<0.001 vs WT mice.

### Overexpression of *Tmem18* within the PVN reduces food intake and increased energy expenditure

Having determined that *Tmem18* expression in the PVN was altered by changes in nutritional state and that global loss of *Tmem18* can affect food intake and energy expenditure, we used stereotactic injection of adeno-associated viral vector (AAV-T18) to manipulate expression of *Tmem18* within the PVN and study subsequent effect on feeding behaviour and body weight. We first undertook targeting studies using AAV over-expressing GFP in a cohort of 12 week old C57/BL6 male mice. Two weeks after unilateral injection, GFP expression was clearly visible within the targeted PVN (Fig 4A). Using identical serotype, volume and co-ordinates as the AAV-GFP experiments, we then performed bilateral injections of an AAV-Tmem18-cDNA into the PVN of 12 week old C57/BL6 male mice to increase *Tmem18* expression specifically within this region of the hypothalamus. At the end of the experiment, mice were sacrificed and brains dissected. Micro-punches of the PVN were obtained from each brain and RNA extracted, cDNA synthesised and Q-RT-PCR performed as detailed in the methods section. Compared to control mice, mice who received vectors expressing *Tmem18* cDNA had a significant two-fold increase in expression level (Fig 4B).

**Figure 4.**
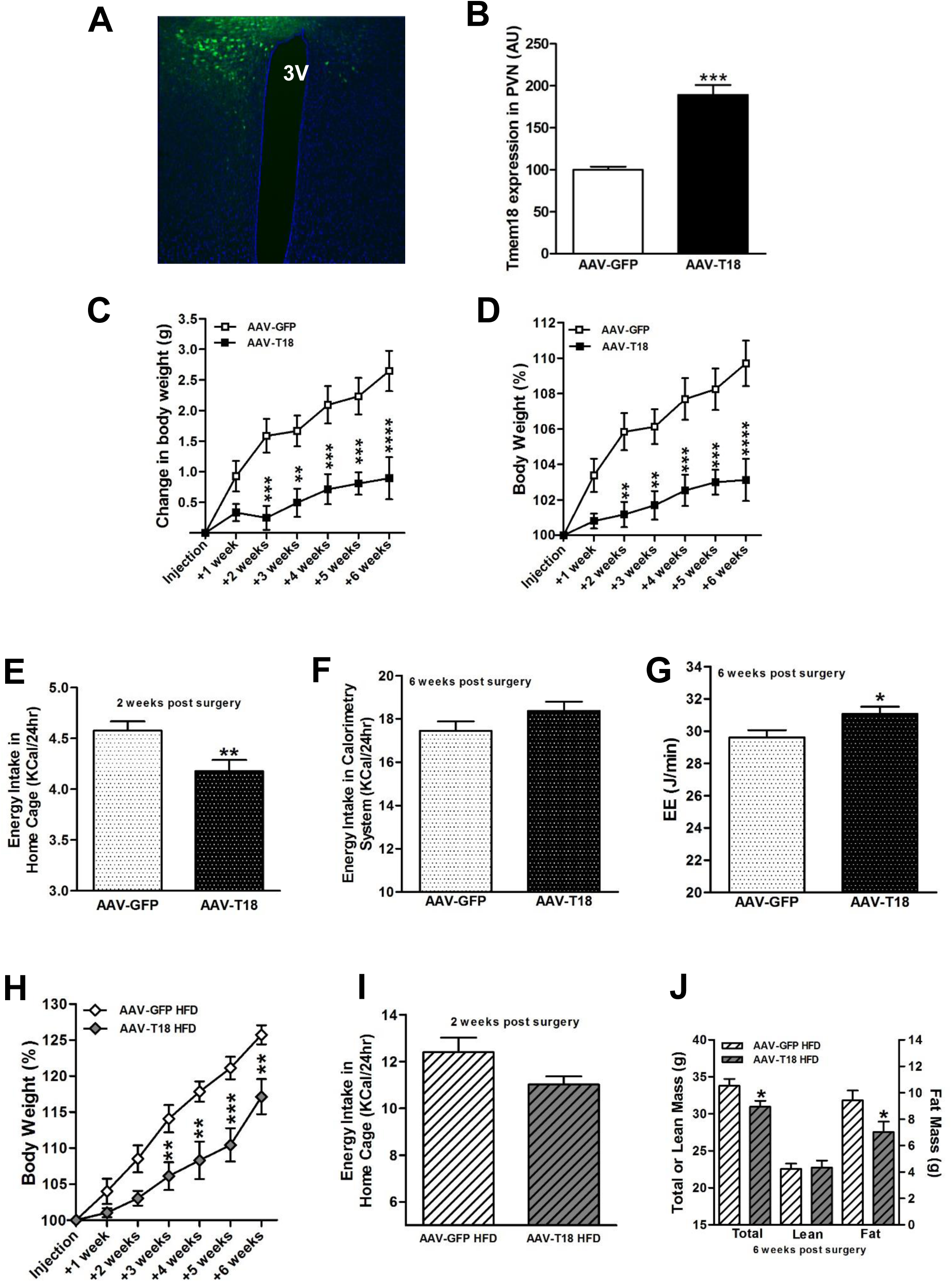
Overexpression of *Tmem18* within the PVN. **(A)** Representative section showing discrete GFP expression (green) within the PVN two weeks after unilateral AAV-GFP injection, with DAPI staining shown in blue. 3V, 3^rd^ ventricle. **(B)** Change in *Tmem18* gene expression within the PVN of mice 7 weeks after bilateral injection of an adeno-asssociated-vector (AAV-T18). **(C)** Change in body weight of mice, measured weekly for 6 weeks after bilateral PVN injections with either an AAV-T18 cDNA or AAV-GFP (n=15 each group). **(D)** Percentage change in body weight of mice, measured weekly for 6 weeks after bi-lateral PVN injections with either an AAV-T18 cDNA or AAV-GFP (n=13 each group). **(E)** Average 24-hour energy intake of mice, measured in their home cage, 2 weeks post-surgery (n=15 each group). **(F)** Average 24-hour energy intake of mice, in the indirect calorimetry cage, 6 weeks post-surgery (n=13 each group). **(G)** ANCOVA analysis of total energy expenditure (EE), assessed over a 24-hour period 6-weeks post injection, with covariates evaluated at body weight=29.41g (n=13 each group). **(H)** Percentage change in body weight of mice placed on a high fat diet immediately after surgery and measured weekly for 6 weeks after bi-lateral PVN injections with either an AAV-T18 cDNA or AAV-GFP (n= 6 each group). **(I)** Average 24-hour energy intake of mice on HFD, measured in their home cage, 2 weeks post-surgery (n= 6 each group). **(J)** Body composition analyses of mice on HFD showing total, lean or fat mass at 6 weeks post surgery (n= 6 each group). AAV-GFP mice are represented by white symbols and bars. AAV-T18 mice are represented by black symbols and bars. AAV-GFP mice on HFD post surgery are represented by white symbols and bars. AAV-T18 mice on HFD post surgery are represented by grey symbols and bars. Data are expressed as mean ± SEM. Time-course data were analysed using the repeated measures ANOVA model. All other p-values were calculated using a two-tailed distribution unpaired Student’s t-test and established statistical significance as follows * p<0.05, ** p<0.01, *** p<0.001 vs GFP injected mice.

Our initial experiments sought to determine what impact overexpression of *Tmem18* within the PVN would have upon energy balance so we followed a cohort of mice for 6 weeks after surgery, recording body weight weekly, measuring food intake in home cage at 2 weeks post-surgery and finally assessing both food intake and energy expenditure at the end of the 6 week period. In terms of absolute change in body weight, there was a significant difference in body weight gain at 2 weeks post-surgery (Fig 4C). By 6 weeks, mean weight gain in mice overexpressing *Tmem18* was only 0.9 g compared to a 2.7 g in the control group injected with an AAV-GFP (Fig 4C). Expressed as a percentage weight gain from starting weight a post-surgery difference between groups was also evident at 2 weeks and remained statistically significant for the remainder of the study period (Fig 4D).

Analysis of food intake at 2 weeks post-surgery demonstrated AAV-T18 treated mice to have a significant reduction in food intake which was likely to have contributed to their reduction in weight gain (Fig 4E). Interestingly, when re-assessed at 6 weeks, this relative hypophagia appeared to have resolved with no difference in energy intake between genotypes (Fig 4F). However, at 6 weeks post-surgery there was a significant difference in energy expenditure with AAV-T18 treated mice showing an increase compared to the control group (Fig 4G, ANCOVA analysis with correction for body weight differences, *p=0.03).

Additionally, another group of mice (n=6) over-expressing *Tmem18* were placed on a high fat diet immediately after stereotactic surgery. Again, in comparison to sham treated animals, mice over-expressing *Tmem18* gained significantly less weight, with this difference becoming significant at three weeks post injection (Fig 4H). There was a trend for a reduction in energy intake at 2 weeks (Fig 4I) and DEXA scanning at 6-weeks post-surgery showed a significant difference in both total and fat mass between the two groups of mice (Fig 4J).

### TMEM18 contains four, not three, transmembrane domains

TMEM18 has been described to be a three transmembrane protein that localizes to the nuclear membrane, courtesy of a nuclear localization signal, and may be involved in transcriptional repression (19, 20). To test this hypothesis, we first embarked on a deep phylogenetic analysis exploring sequence profile-to-profile homology (28) with MPI Bioinformatics Toolkit software (29). Database searches revealed remote but clear homology of TMEM18 to various ion channels. These included proteins from Pfam families (http://pfam.xfam.org) of fungal transient receptor potential ion channels (PF06011) and bacterial mechanosensitive ion channels (PF12794). Searches in metazoan proteomes yielded Unc-93 A homologues (mammalian gene UNC93A) annotated in *C. elegans* as an ion channel. The top hits (Toolkit parameter of probability of true positives about 80%) comprised a voltage-gated sodium channel from *Caldalkalibacillus thermarum* and an ion transport protein from *Arcobacter butzleri* (PDB identifiers 4bgn and 3rvy). Although their mutual phylogenetic distances are remote, they fold into a very similar structure and TMEM18 shares suggestive homology with their ion-transmitting domain. This region forms an oligomer consisting of two N- and C-terminal transmembrane helices pointing to the cytoplasm and two shorter inner helices fully embedded in the membrane. The conserved charged tip of their C-terminal helices is entirely outside the membrane and participates in the pore opening. It is highly probable that TMEM18 possesses a very similar topology and structure. Based on the homology to the *C. thermarum* channel, a very tentative model can be proposed for further testing. Figure 5A shows the modelled membrane topology and a putative 3D structure under the assumption that TMEM18 forms a tetramer similar to the *A. butzleri* channel (30). As a nuclear membrane protein, it should expose both its termini to the cytoplasm and therefore would be comprised of four, rather than three, transmembrane domains. To test this experimentally we transfected Cos cells with a N-terminal FLAG-tagged *Tmem18* construct then detected protein expression using either a FLAG antibody (to identify the N-terminus of the protein) or a TMEM18 antibody raised against C-terminus amino acids 120-134 (to identify the C-terminus of the protein). Protein expression was analyzed using two different detergents, TX100 that permeabilises all cellular membranes and digitonin, that only permeabilises the plasma membrane at a concentration of 40ug/ml. Control antibodies (lamin B and calnexin) gave the expected results. Lamin B, which is localized to the inside of the nuclear membrane could only be detected when TX100 is used to permeabilise both the plasma and nuclear membrane (Fig 5B & 5C) whereas calnexin, which spans the nuclear membrane with the C-terminus pointing into the cytoplasm, could be detected with both permeabilisation reagents using a C-terminal antibody (Fig 5D & 5E). Both the N-terminus (FLAG antibody, Fig 5F & 5G) and C-terminus (TMEM18 antibody, Fig 5H & 5I) of TMEM18 could be detected with digitonin permeabilisation indicating that both ends of the protein were located within the cytoplasm and confirming the bioinformatics analysis which suggested that Tmem18 is a four transmembrane protein.

**Figure 5.**
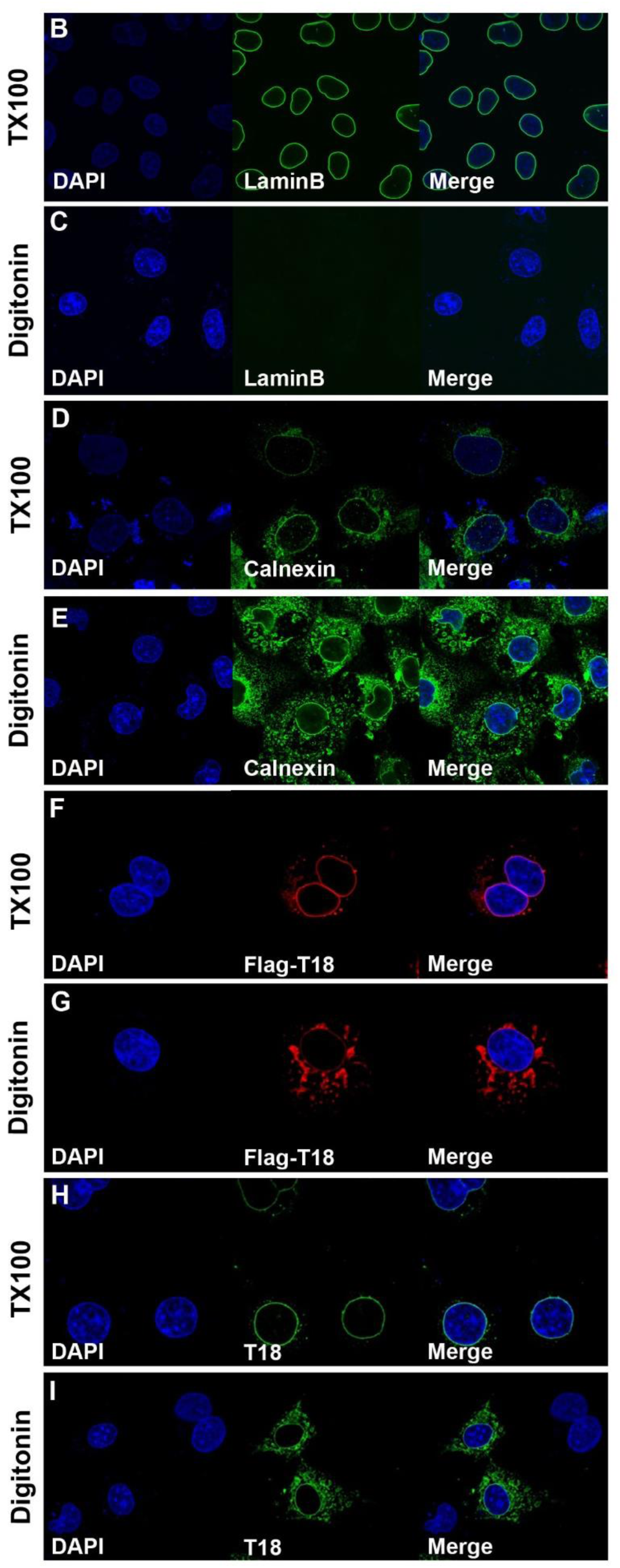
Topology of the TMEM18 protein. **(A)** Tentative illustrative sketch of TMEM18 modelled as a tetramer. The individual monomers are displayed in different colours. The conserved C-terminus (∼25 amino acids) is exposed to the cytoplasm and the upper four-helix domain is embedded in the membrane with its N-terminus pointing to the cytoplasm. The preceding sequence of ∼50 amino acids is exposed to the cytoplasm but cannot be modelled due to the lack of homology. **(B-H)** Overexpression of N-terminal FLAG-tagged TMEM18 in COS cells treated with either TX-100 (permeabilises both plasma & nuclear membrane) or digitonin (permeabilises plasma membrane only). Lamin **(B&C)** and Calnexin **(D&E)** are shown as controls. TMEM18 expression was detected with either a FLAG antibody **(red, F&G)** or an antibody to the C-terminus of TMEM18 **(green, H&I)**.

### TMEM18 is unlikely to directly regulate transcription

Little is known about the function of TMEM18. However, it has been suggested that it binds DNA and supresses transcription (20). To test this hypothesis, we performed RNA-Seq analysis of hypothalami from wildtype and *Tmem18* knockout male mice (ENA accession PRJEB13884).

Altogether, 27691 (27727 with non-zero total read count; 36 outliers per cooks Cutoff (31)) annotated genes had reads mapped in at least one sample and were used for differential expression analysis (average number of uniquely mapped reads per sample: 32.4×10-6 ± 2.04×10-6 SD). Interestingly, after adjustment for multiple testing, only *Tmem18* was significantly differentially expressed within the hypothalami of the two genotypes (padj = 3.02 × 10-22, Fig 6A). Q RT PCR analysis of *Tmem18*, and six additional genes with smallest (but not statistically significant) padj values confirmed the RNAseq data (Fig 6A and 6B), suggesting that TMEM18 is unlikely to be a global regulator of transcription as previously reported (20).

**Figure 6.**
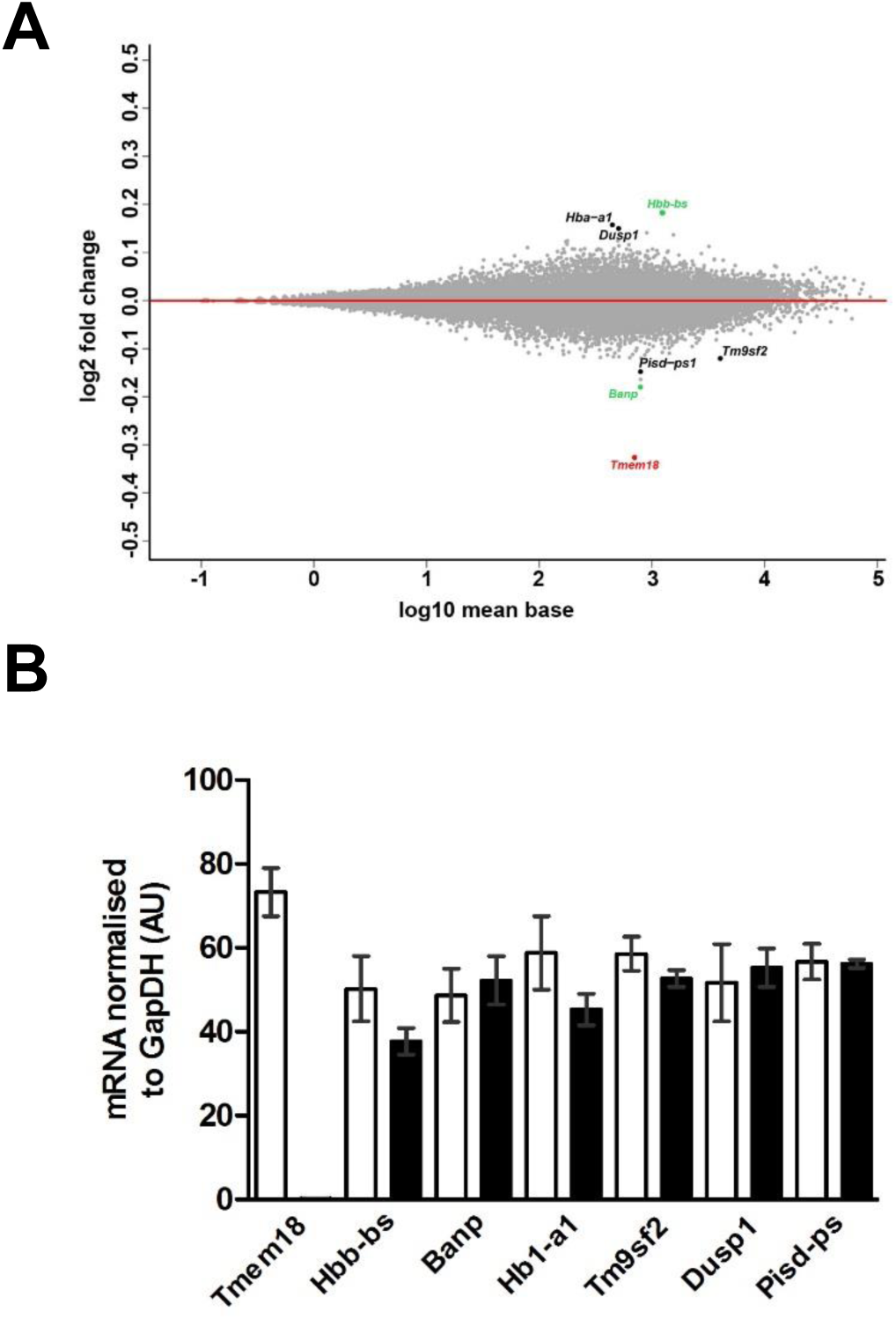
Effect of loss of *Tmem18* on the hypothalamic transcriptome. **(A)** MA plot showing differentially expressed genes in the hypothalami of *Tmem18^wt/wt^* and *Tmem18^tm1a/tm1a^* male mice. *Tmem18*, the only gene with a statistically significant difference in expression, is shown in red. The next two transcripts with the largest difference in expression (*Hbb-bs* and *Banp*) are shown in green. Four additional genes selected for confirmatory analysis are shown in black. **(B)** Q-RT-PCR analysis of selected genes in the hypothalami of *Tmem18^wt/wt^* and *Tmem18^tm1a/tm1a^* male mice.

### TMEM18 interacts with two nuclear pore proteins, NDC1 and AAASs

We used the methods of affinity purification and mass spectrometry (32) to identify novel TMEM18 interacting partners. Either FLAG-tagged TMEM18 or empty FLAG vector were over expressed in HEK293 cells. Protein complexes were immunoprecipitated using a FLAG antibody and mass spectrometry analysis performed to detect proteins bound to TMEM18. This process identified 116 proteins pulled down in control cells expressing the empty FLAG vector, 221 proteins common to both the control and experimental group, and 244 proteins unique to cells expressing FLAG-TMEM18 (See Supplementary Table 1 for a list of all proteins identified as being pulled down by FLAG-TMEM18). Interestingly, three members of the nuclear pore complex, NDC1, AAAS, and NUP35/53 (33) showed high numbers of assigned spectra indicating a high abundance of these proteins after pulldown. Therefore, biomolecular immunofluorescence complementation (BiFC) assays were employed to confirm interactions between these proteins and Tmem18. Controls behaved as expected and showed that no YFP expression was seen if YN constructs were co-transfected with a YC-STOP plasmid (Fig 7A). YN-FLAG-NDC1 and YC-AAAS served as an appropriate positive control (Fig 7B) as these proteins have been reported to interact previously (34, 35). YFP expression could be detected in cells expressing YC-TMEM18 and either YN-FLAG-NDC1 or YN-FLAG-AAAS; but not YN-FLAG-NUP35 (Fig 7C-G). Positive BiFC protein-protein interactions were corroborated further by co-immunoprecipitation experiments using FLAG-TMEM18 and either GFP tagged NDC1 (Fig 7H) or GFP tagged AAAS (Fig 7I).

**Figure 7.**
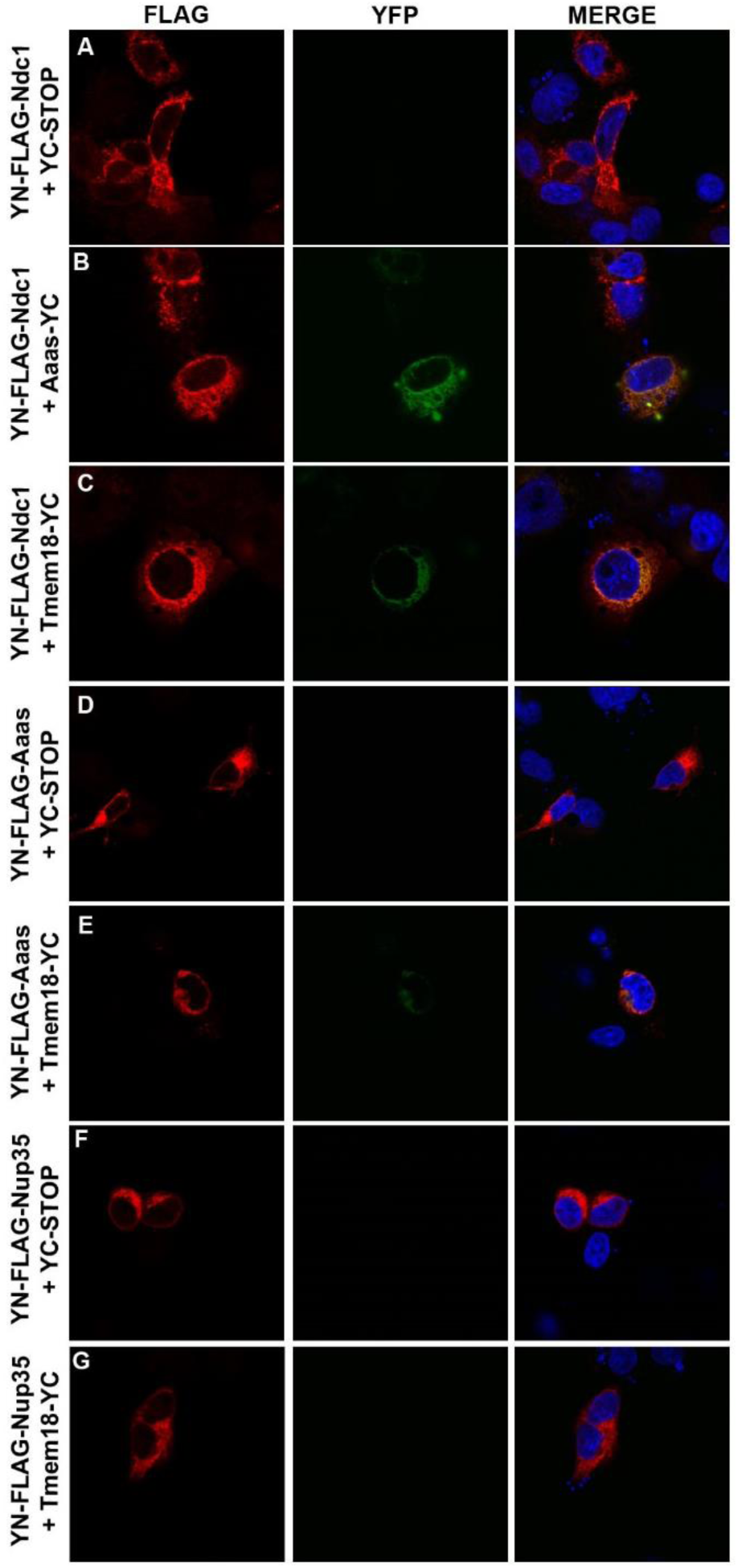
BiFC and co-IP confirmation of Tmem18 interaction with Ndc1 and Aaas. **(A-G)** Physical interaction between Tmem18 and Ndc1 and Aaas was confirmed by BiFC. The N-terminus of YFP was fused to FLAG-tagged Ndc1 (YN-FLAG-Ndc1, panel C) or Aaas (YN-FLAG-Aaas, panel E) or Nup35 (negative control, YN-FLAG-Nup35, panel G), while the C-terminus of YFP was fused to Tmem18 (Tmem18-Yc). FLAG expression was detected using a FLAG antibody (red) while YFP signal is depicted in green. YN-FLAG-Ndc1 and Aaas-YC was used as a positive BiFC control (panel B). **(H and I)** Co-immunoprecipitation experiments using FLAG-tagged Tmem18 and GFP-tagged Ndc1 **(H)** or Aaas **(I)** overexpressed in HEK cells. Dashed lines in the blot represents two different lanes in the same blot put together.

## DISCUSSION

Our data indicate that altering *Tmem18* expression in mice, both globally and within the hypothalamus, can alter body weight. Male mice with a germline loss of *Tmem18* have an increased body weight due to a significant increase in both fat and lean mass. This phenotype is more pronounced on a high fat diet, where weight gain is driven by hyperphagia. In contrast, overexpression of *Tmem18* expression within the hypothalamic PVN can reduce food intake, increase energy expenditure and reduce both total body and fat mass.

The increased body weight phenotype of *Tmem18* null mice was sexually dimorphic. Sex-specific differences in metabolic studies focusing on body weight are not uncommon and have been seen, for example, after embryonic TrkB inhibition or with disruption of GABA receptor signalling in proopiomelanocortin neurons (36, 37). In these reports and in our current study data, the mechanisms behind the sex-specific changes in body weight remain to be fully determined but the differences in circulating gonadal derived hormones remain, of course, a potential contributor. Indeed, the relevance of oestrogen in influencing the response to a dietary intervention was highlighted again in a study by Dakin *et. al*. in which oestradiol treatment to male mice fed a high-fat, high-sugar ‘obesogenic’ diet prevented increases in adipose tissue mass (38).

Male mice on a high fat diet gained weight because of increased energy intake. Of note, data from indirect calorimetry also showed *Tmem18* null mice to have a ∼10% increase in energy expenditure compared to wild type mice, but as food intake was increased by ∼30% the dominant drive was to caloric excess and weight gain. This could be considered an example of “diet induced thermogenesis”, a state seen as an effect of a change to a higher calorie diet in which mice increase energy expenditure and food intake simultaneously (39). The potential biological purpose and the tissues relevant to this phenomenon continue to be disputed (40). There is more agreement that the ambient temperature of a study can have a strongly qualitative effect on the outcome of metabolic studies (41). Our studies were conducted under “standard” animal house conditions that can be considered a chronic thermal stress to mice. Future studies of energy expenditure in *Tmem18* deficient mice living at thermoneutrality may be helpful in further understanding the role of this molecule in metabolic control.

When studying high fat fed male animals, in both genotypes food intake was higher in the home cage compared to that recorded in the calorimetry system. The difference in absolute food intake may reflect a degree of stress in a less familiar environment but we mitigated against this by allowing mice a period of acclimatisation in both the room and the cage type in which the calorimetry studies were performed.

In trying to determine the locus of action driving the effects of TMEM18 on body composition we chose to start within the PVN, an anatomical region enriched with neuronal populations involved in appetitive behaviour and energy expenditure (42-46). In two distinct cohorts, one fed normal chow, another high fat diet, adult onset overexpression of *Tmem18* in the PVN diminished body weight gain. There was also a reduction in food intake and when viewed with the hyperphagia seen in the *Tmem18* null mice, our data suggest that *Tmem18* expressing neurons in the PVN have a role in food intake. However, our data also indicate that *Tmem18* is expressed within a number of other regions within the hypothalamus. Further the nature of the different perturbations (germline loss in *Tmem18^tm1a^ vs* delivery of AAV via stereotactic injection) means that the PVN neuronal population which lost *Tmem18* expression in the null mice may not be the same population that gained overexpression in the stereotactic-driven AAV study. We were unfortunately unsuccessful in our attempts to use siRNA and *Cre* to selectively knock down and/or delete *Tmem18* in the PVN despite promising preliminary studies in vitro (data not shown). In the future, transgenic lines with specific anatomical *Cre* drivers (e.g. *Sim1-Cre, Agrp-Cre*) may be helpful to further delineate the role of Tmem18 in hypothalamic nuclei.

In recent years there have been several reports investigating *Tmem18* expression within the brain and its link to obesity. Almen *et al*. reported that *Tmem18* is expressed in tissues throughout the body and confirmed abundant *Tmem18* expression in the hypothalamus (19). They measured expression across a range of different nutritional states (48-hour consumption of sucrose or intralipid, 16 hour fast and 24 hour fast) but *Tmem18* mRNA levels in whole hypothalami remained unchanged. Yoganathan *et al*. also examined expression of *Tmem18* in whole hypothalami from fasted mice (4 hours only) and from mice fed both standard chow and a high sucrose diet (47). Again, expression levels in the hypothalamus were identical across these conditions although intriguingly in the rest of the brain excluding the hypothalamus, *Tmem18* expression was some 8-fold higher in the fed group compared to the fasted group. Schmid *et al*. studied *Tmem18* expression in obese Zucker diabetic fatty rats and again found the highest amounts of *Tmem18* mRNA in the hypothalamus (48). Thus, although several groups have also shown *Tmem18* to be highly expressed within the hypothalamus, not all have reported it to be nutritionally regulated. However, the specific neuronal populations within the relatively heterogeneous PVN through which *Tmem18* functions remain to be fully defined.

Our report presents interesting new data on the potential function of TMEM18. Previous reports have indicated that TMEM18 is highly conserved, has no family members, localizes at the nuclear membrane and that the C-terminus of the protein may be involved in the regulation of transcription. Whilst the localization of TMEM18 is not disputed we propose that a role for TMEM18 in transcriptional regulation is unlikely given our RNA-Seq data showing that the hypothalamic transcriptome of *Tmem18^tm1a^* mice is not significantly different to that of wildtype littermates. Previous bioinformatic analysis had predicted a 3 transmembrane structure (19, 49). However, our cellular experiments reveal that TMEM18 is in fact comprised of four transmembrane segments indicating that it is unlikely that the C-terminus is involved in binding DNA as earlier reported (20). Instead we hypothesise that TMEM18 may be involved in the transport of molecules across the nuclear envelope. Pulldown experiments, corroborated by BiFC and IP studies have identified two novel binding partners for TMEM18 – NDC1 and AAAS/ALADIN. These proteins are two of the thirty NUPs that make up the nuclear pore complex (NPC) (50). As NDC1 and AAAS have previously been reported to interact with one another (35) it is possible that TMEM18 is not interacting directly with both proteins. By enabling transport of molecules across the nuclear envelope, NUPs are involved in many fundamental cellular processes, consequently abnormal expression/function of these NUPs has been linked to various human diseases such as cancer, cardiovascular disorders, autoimmune disease and neurological defects (51). Further investigation into the interaction of TMEM18 with members of the NPC will hopefully shed new light on the molecular function of TMEM18 and its role in obesity.

Prior to featuring in any GWAS, *Tmem18* was initially identified in a bioinformatics screen investigating *cis*-regulatory motifs involved in translational control (52) and an additional screen searching for novel modulators of cell migration (49). The association of single nucleotide polymorphisms (SNPs) near human *TMEM18* with obesity was first reported by Willer *et al*. (2) and has since been identified by multiple GWAS (reviewed in (6)). Undoubtedly, GWAS have produced much new data relevant to obesity. However, increasing knowledge about the association between genetic variants and the condition under investigation brings forth a series of challenges. The GWAS data alone cannot indicate which SNPs are causal nor which cell type or tissue is relevant. Further, many of the variants are located in non-coding regions of the genome giving no immediate clue to mechanism. All of these points are germane to the SNPs located near to *TMEM18*. SNP rs6548238 is, for example, greater than 30 kb downstream of *TMEM18* and falls in a relative ‘gene desert’, lying over 300 kb from the next nearest genes (*SNTG2* and *FAM150B*). Thus, although these loci have taken on the moniker of ‘near to *TMEM18*’, to date it has remained unclear whether these variants have any influence on the regulation of *TMEM18* expression or function, or indeed if TMEM18 has a direct role in energy homeostasis.

Efforts to advance understanding from association to biologically relevant mechanisms have combined multiple techniques and evidence from several model platforms. For example, when investigating obesity associated noncoding sequences within *FTO*, Smemo used chromatin conformation capture techniques (CCST) to show a region of *FTO* directly interacts with, and forms part of the regulatory landscape of, the homeobox gene *IRX3* (10). They also reported that relevant SNPS were associated with expression of *IRX3*, but not *FTO*, in human brains. In a more recent study of obesity associated *FTO* variants, Claussnitzer combined chromatin capture technology, eQTL data and a powerful bioinformatics approach to report that these variants can reduce thermogenesis in human adipocytes (53). However, given the clear evidence that the obesogenic *FTO* SNPs have a stringent and reproducible impact on aspects of human appetite (54) including directly measured food intake (55) coupled with the lack of evidence for any impact on human energy expenditure (56), the physiological relevance of these latter observations must remain open to question.

Thus far our attempts to interrogate the region near *TMEM18* through the examination of various gene expression databases shows no eQTL data relevant to these SNPs (data not shown). A brain tissue specific approach may prove more fruitful but unfortunately RNA expression data specifically from human hypothalamus linked to genotype is not a currently publicly available resource.

The studies we have described have strengthened the candidacy of *TMEM18* as, at least in part, the mediator of the association between genetic variation in this region of chromosome 2 and human adiposity, one of the strongest associations of a common variant with human obesity. Our work has also provided novel information about the topology and binding partners of this enigmatic protein associated with the nuclear pore complex.

## METHODS

### Animals

All procedures were carried out in accordance with guidelines of the United Kingdom Home Office. Animals were kept under controlled temperature (22^°^C) and a 12-h light, 12-h dark schedule (lights on 7:00–19:00). Standard chow (Special Diet Services, SDS) or a 45% fat diet (D12451, SDS) and water were available *ad libitum*. Mice carrying the knockout first conditional-ready allele *Tmem18^tm1a^*^(^*^EUCOMM^*^)^*^Wtsi^*, abbreviated to Tmem18^tm1a^ in this paper, were generated on a C57BL/6N background as part of the Sanger Mouse Genetics Project (MGP) for the European Conditional Mouse Mutagenesis Program [EUCOMM]. Detailed description of the Sanger Mouse Genetics Project methodology has previously been reported (23, 57-59). Germ line transmission was confirmed by a series of genotyping PCR analyses (http://www.knockoutmouse.org/kb/25/). Following confirmation, mice derived from heterozygous inter cross, were genotyped for the *Tmem18^tm1a^* allele by PCR. *Tmem18^flox/flox^* mice (Tm1c line) were generated by crossing Tmem18^*tm1a*^ mice to a Flpase deleter line on the same genetic background (60).

### Design and Construction of AAV Vectors

To generate a *Tmem18* expressing adeno associated viral (AAV) vector, the cDNA of murine *Tmem18* was cloned into an AAV backbone plasmid under the control of the CMV promoter. AAV vectors expressing GFP under the control of the same promoter were used in a control group of animals as well as to test for optimum titre and accuracy of intranuclear injections in preliminary experiments.

### Viral production and purification

AAV vectors were generated by helper virus-free transfection of HEK293 cells using three plasmids. Vectors were purified by two consecutives caesium chloride gradients using an optimized method (61), dialyzed against PBS, filtered, titred by qPCR and stored at −80^°^C until use. This study used AAV serotype 7.

### Stereotactic surgery

Mice were stereotaxically injected with AAV while under isoflurane induced anaesthesia. The coordinates used for the injections were 1.0 mm caudal to bregma, ±0.25 mm lateral to the midline and 5.0 mm below the surface of the skull in all cases. Using a 10 μl Hamilton syringe and a 33 gauge needle, 200nl of AAV solution was injected into each side of the hypothalamus over a 1 minute period. After delivery of the virus the needle was left in place for 10 minutes to prevent reflux. The titre of the vector preparations were as follows: AAV7-CMV-GFP 2.1×10^12^vg/ml (dose delivered per 200 nl 4×10^7^vg); AAV7-CMV-Tmem18 6.3×10^12^vg/ml (dose delivered per 200 nl 1.2×10^8^vg).

### Metabolic phenotyping

Body weight measurements were recorded on a weekly basis. Food intake studies were carried out on single housed animals in home cages. Daily food intake was an average taken from 10 consecutive days. Body composition was determined using dual-energy x-ray absorptiometry (DEXA) (Lunar PIXImus2 mouse densitometer, General Electric Medical Systems, Fitchburg, WI). Any animal who did not remain weight stable did not proceed to calorimetry. Energy expenditure was determined using indirect calorimetry in a custom built monitoring system at 20^°^C (Ideas Studio, Cambridge, UK). Activity was assessed by beam breaks. Beams were 1.25 cm apart and activity measurements were taken to be total beam breaks, rather than consecutive beam breaks. Any data from an animal whose weight changes in the calorimetry system were beyond those seen in the long-term home cage environment were excluded from analysis. At the end of the experiment animals were sacrificed and tissues collected for further analysis.

### Immunohistochemistry and histology

Animals were anesthetized with pentabarbitol (Dolethal) then transcardially perfused with 20 ml PBS followed by 40 ml 10% Formalin/PBS (Sigma). Brains were dissected out and incubated in 15% sucrose/10% Formalin overnight at 4^°^C. Following cryoprotection in 30% sucrose/PBS, brains were frozen on dry-ice and stored at −80^°^C overnight. Serial 35 μΜ sections were taken using a freezing microtome and mounted on glass slides (VWR).

### Laser capture microdissection and Q-RT-PCR of fed and 48hr fasted mouse brains

Laser-captured microdissection and total RNA isolation were carried out as previously described (22, 62).

### Laser capture microdissection and RNA Sequencing of fed and fasted mouse brains

Coronal sections of 20 um thickness were prepared on a cryostat, mounted on RNase-free membrane-coated glass slides with laser microdissection performed using a P.A.L.M. MicrolaserSystem (P.A.L.M. Microlaser Technologies). 2ng of total RNA from each sample was SPIA (single primer isothermal amplification) amplified with an Ovation RNA-Seq System V2 (NuGEN), after which cDNA libraries were prepared using the Encore Rapid DR Multiplex System (NuGEN). Single-end reads (SE50) were sequenced on an Illumina Hi-Seq 2500. Reads were mapped to *Mus Musculus* GRCm38 genome assembly (Ensembl) with Tophat, and comparisons of fasted vs fed samples across the four hypothalamic were performed in RStudio using the edgeR and limma packages.

### RNA-Seq analysis of whole hypothalami from wildtype and Tmem18^tm1a/tm1a^ mice

The hypothalamus was dissected from 4 male adult mutant mice and 5 wild-type littermate controls. RNA was extracted using Qiagen RNeasy Mini Kit according to manufacturer’s instructions. 1 ug of RNA was used to prepare multiplexed sequencing libraries with an Illumina RNA Library Preparation Kits as per the manufacturer’s instructions. Paired end sequencing was performed on the Illumina Hi-Seq 2500, generating 75 bp, stranded paired-end reads. Using STAR (v.2.4) (63) the sequenced reads were aligned to a modified version of the mouse reference genome (GRCm38) with addition of pseudo-chromosomes containing sequences of the knockout first conditional-ready allele (specifically, of the LacZ and neomycin regions of the cassette) for genotyping quality control (64). The number of reads that mapped to each annotated gene was counted using the HTSeq (v.0.6.1) count function (HTSeq.scripts.count, mode *intersection-nonempty*). The DESeq2 package (v.1.9.29) (31) was used for differential gene expression analysis between mutant and wildtype mice, using the Benjamini-Hochberg procedure (to control for multiple testing, returning an adjusted p value (padj)). A significance cut-off of padj < 0.05 was used.

### Quantitative-RT-PCR

Q-RT-PCR was performed on whole hypothalami and brown adipose tissue (BAT). Tissues were dissected from individual mice, snap frozen and stored at -80^°^C until processed. RNA was extracted using TriSure (Bioline) and reverse transcribed using Il-Prom-II reverse transcription system (Promega) according to manufacturer’s instructions. To assess Tmem18 expression within the PVN following surgery micro-punches of the PVN from individual mice were obtained from fresh frozen brains. Accuracy of the micro-punches was confirmed with cresyl violet staining of the cryostat sections after nuclei removal. RNA was isolated from the micro-punch using the Qiagen RNeasy micro system and 50 ng of purified RNA were used in a random-primed first strand cDNA synthesis reaction, using the VILO reverse transcriptase kit (Invitrogen). Quantitative PCR reactions were performed in triplicate on an ABI 7900HT (Applied Biosystems) using ABI PCR master mix and commercially available Taqman probes, according to manufacturer’s protocols. All Q-RT-PCR data was normalised to *Gapdh* expression.

### Transient transfection and immunocytochemistry

Cos7 cells were plated on four-well glass chamber slides (Nalge Nunc International) at 25,000 cells/ml; 16 h later, each well was transfected with 500 ng of plasmid DNA using Lipofectamine (Thermo Fisher Scientific). After 48 h at 37^°^C, cells were fixed in 3.7% formaldehyde with either 0.2% TritonX100 for 10 mins at RT or 40 ug/ml Digitonin for 10 mins at 4^°^C. After washing in PBS, cells were incubated O/N at 4^°^C with the appropriate primary antibody. The following morning, cells were washed in PBS and incubated with an appropriate secondary antibody conjugated to either a 488 or 594 fluorophore. After washing in PBS, coverslips were mounted using VectaShield hardset mounting media with DAPI (Vector Laboratories) and fluorescent cells detected using a Nikon Eclipse TE2000-U microscope. If colocalization of two proteins was being studied, before mounting with coverslips, cells were incubated with the second primary antibody at RT for 3hrs, washed in PBS then incubated with an appropriate secondary antibody conjugated to either a 488 or 594 fluorophore. Primary antibodies used were as follows: custom made rabbit-anti-mouseTmem18 antibody raised against amino acids 120-134; Rabbit Polyclonal FLAG tag (Sigma); Monoclonal Anti-FLAG M2 (Sigma), Rabbit-anti-human calnexin (ab22595, Abcam); Goat-anti-mouse lamin B (Santa Cruz Biotech, sc-6217) and Rabbit polyclonal Myc tag (Abcam, ab9106).

### Pulldown and Mass Spec analysis

HEK293 cells were cultured in Dulbecco’s Modified Minimal Essential Medium (DMEM) supplemented with 10% FCS at 37oC under 5% CO2. For protein over expression studies, transient transfection in HEK293 cells was performed using a CalPhos kit according to the manufacturer’s protocol. 48 hours post transfection, cells were harvested and Flag immunoprecipitations (IPs) were performed from the resulting lysates as described (65).

For MS analysis, Flag immunoprecipitates were subjected to SDS-PAGE, each lane was excised and cut into 6 chunks with the proteins digested in-gel using trypsin. The resulting tryptic peptides were eluted and analysed by LC-MSMS using an LTQ-OrbiTrap XL (Thermo) coupled to a nanoAcquity (Waters) in a top 6 DDA fashion. Raw files were processed using MaxQuant 1.3.0.5. Searches were performed using the Andromeda search engine against a Uniprot Mus musculus database (downloaded 14/08/12). Oxidation (M), acetylation (protein N-terminus) and deamidation (NQ) were set as variable modifications. Data was filtered to a peptide and protein FDR of 0.01. Validation of MS analysis results was performed as previously described (65).

### Direct and Competitive Bimolecular Fluorescence Complementation

YFP constructs and controls were a kind gift from David Savage (66). Interactions between YN-FLAG-NDC1/AAAS/NUP35 and MYC-TMEM18-YC in COS-7 cells were assessed by transfecting 200 ng of each YN construct with 200 ng of each YC construct. Four hours post transfection, cells were incubated at 32°C for 20hr and 30°C for 2.5hr to promote the formation of YFP. Twenty-four hours post transfection, cells were fixed, permeabilized, and stained with anti-FLAG mouse monoclonal antibody (Sigma) followed by goat anti-mouse Alexa 594 secondary antibody (Abcam).

### Bioinformatic Analysis

TMEM18 orthologues (144 sequences) from OMA database (http://omabrowser.org) were aligned with Muscle (67) and submitted to HHpred searches (28) in all available microbial fungal and metazoan protein databases and collections of HMM profiles using MPI Toolkit suite (29). The homology models were based on the HHpred alignments. The structures were displayed and examined with Protein Workshop Viewer (68).

### Statistical Analysis

All values are expressed as mean ±S.E.M. Statistical analysis was performed using Graph Pad Prism software (GraphPad Prism) or SPSS (IBM). For the analysis of food intake and body weight over time, two-way repeated measures ANOVA, with a Bonferroni post-test, was used with time and treatment as variables for comparison. For energy expenditure (EE) analysis of covariance (ANCOVA) was performed to assess body weight/EE interactions with body weight as a covariate, genotype or AAV treatment as a fixed factor and EE as a dependent variable. Multiple linear regression analysis was carried out with no selection criteria. Significant differences were designated as *p* < 0.05.

## AUTHOR CONTRIBUTIONS

RL, MS, LT, DR, PG, RA, SV, VS, GY and APC conducted the experiments and/or analysed data. JPW and BL conducted and analysed all RNA Seq experiments on fed and fasted wild-type mice. EA and FB produced and purified viral vectors. CD and DWL conducted and analysed all RNA Seq experiments on *Tmem18* knockout tissue. RL, GY, SO’R and APC designed the experiments. RL, SO’R and APC wrote the manuscript.

## ACKNOWLEDGEMENTS

The authors thank Helen Westby, Will Gee and Elizabeth Wynn for technical assistance. We also thank Satish Patel and Koini Lim for assistance with the BiFC protocol. RL, YCLT, DR, GSHY, SOR and APC are funded by the Medical Research Council (MRC) Metabolic Disease Unit (MRC_MC_UU_12012/1) and animal work was carried out with the assistance of MRC Disease Model Core of the Wellcome Trust MRC Institute of Metabolic Sciences (MRC_MC_UU_12012/5 and Wellcome Trust Strategic Award (100574/Z/12/Z). F. Bosch is the recipient of an award from the ICREA Academia, Generalitat de Catalunya, Spain. Vector generation and production were funded by Ministerio de Economía y Competitividad (SAF 2014-54866-R), Spain. CD and DWL were supported by the Wellcome Trust (WT098051) and CD was supported by the Wellcome Trust PhD Programme for Clinicians (100679/Z/12/Z).

---

**Supplementary Figure 1.**
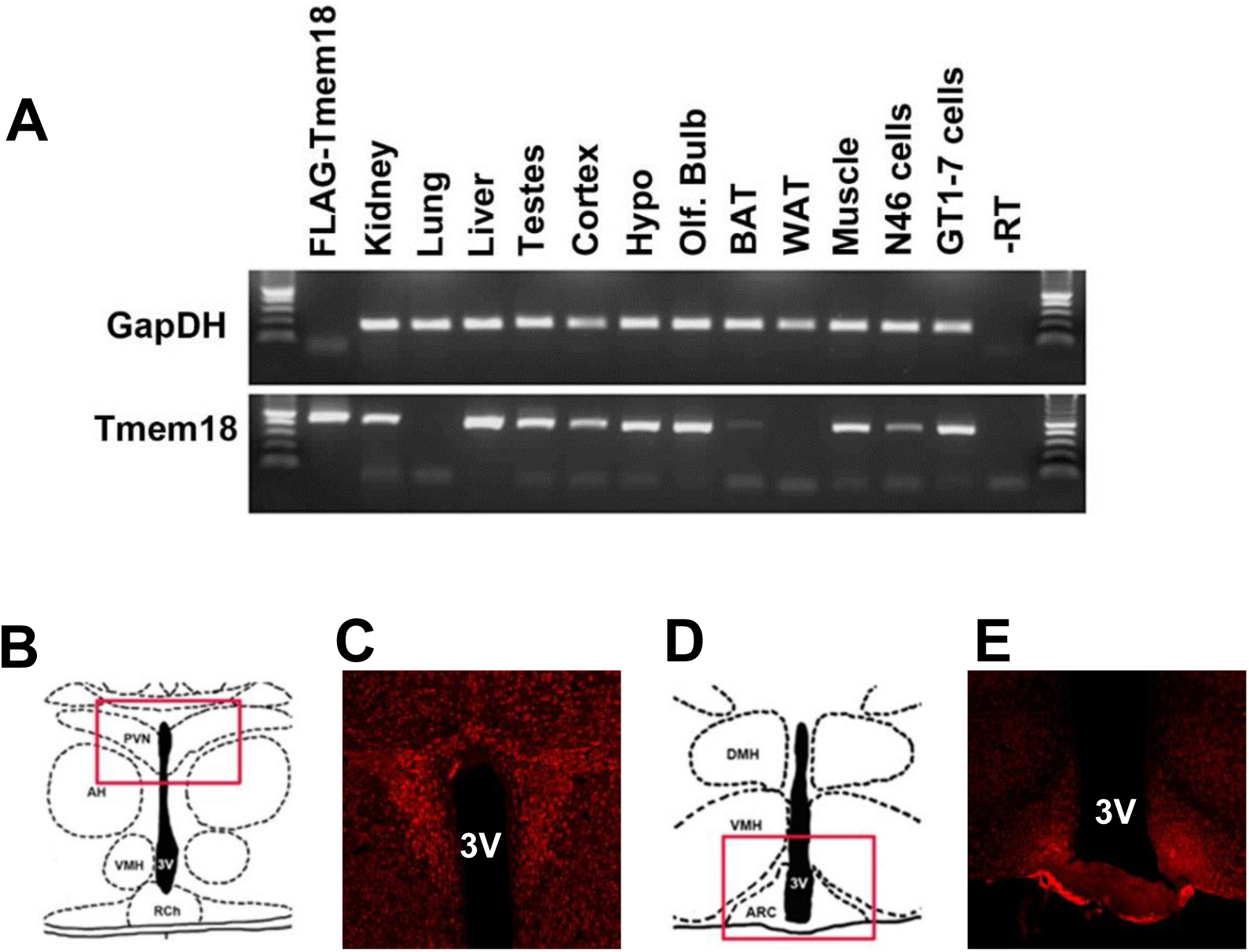
Analysis of *Tmem18* expression. **(A)** RT-PCR analysis of *Tmem18* expression in various mouse tissues. GapDH was used as a positive control. **(B-E)** Schematic showing coronal section through a mouse brain indicating the (**B**) PVN (−1.06 from Bregma) and (**D**) Arc (−1.58 from Bregma). The red boxes indicate the areas in representative image showing immunohistochemical staining for TMEM18 in (**C**) PVN and (**E**) Arc (coronal sections (X10), adult brain). AH; anterior hypothalamus; ARC, arcuate nucleus; DMH, dorsomedial hypothalamus; PVN, paraventriular nucleus; RCh, retrochiasmatic area; VMH, ventromedial hypothalamus; 3V, third ventricle.

**Supplementary Figure 2.**
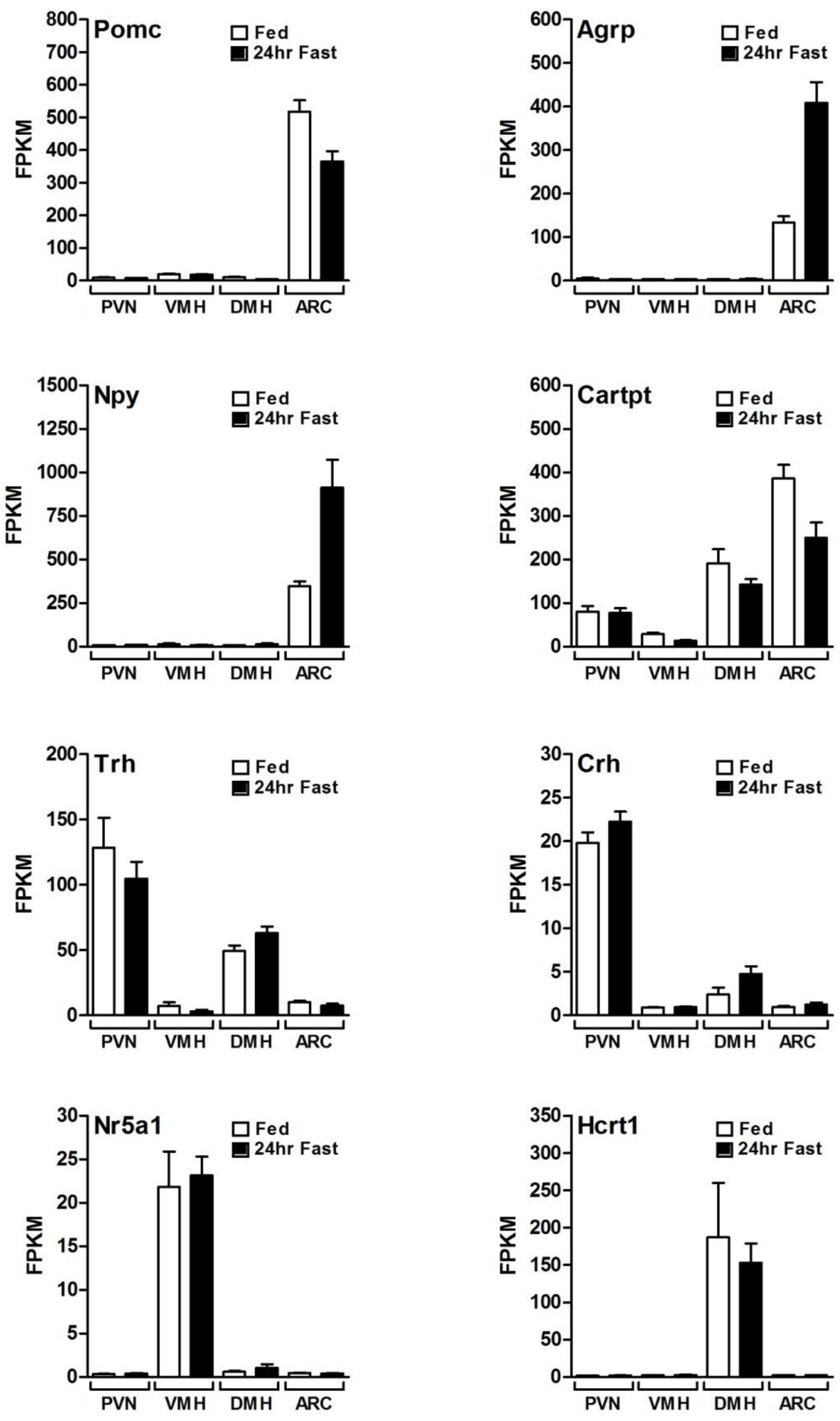
Validation of laser-capture microdissection specificity in *ad libitum fed* and 24hr fasted wildtype mice. Bar charts showing FPKM values for genes known to be enriched in given hypothalamic nuclei. White bars show fed mice, black bars show mice fasted for 24hrs. PVN, paraventricular nucleus; VMH, ventral medial hypothalamus; DMH, dorsal medial hypothalamus; ARC, arcuate nucleus.

**Supplementary Figure 3.**
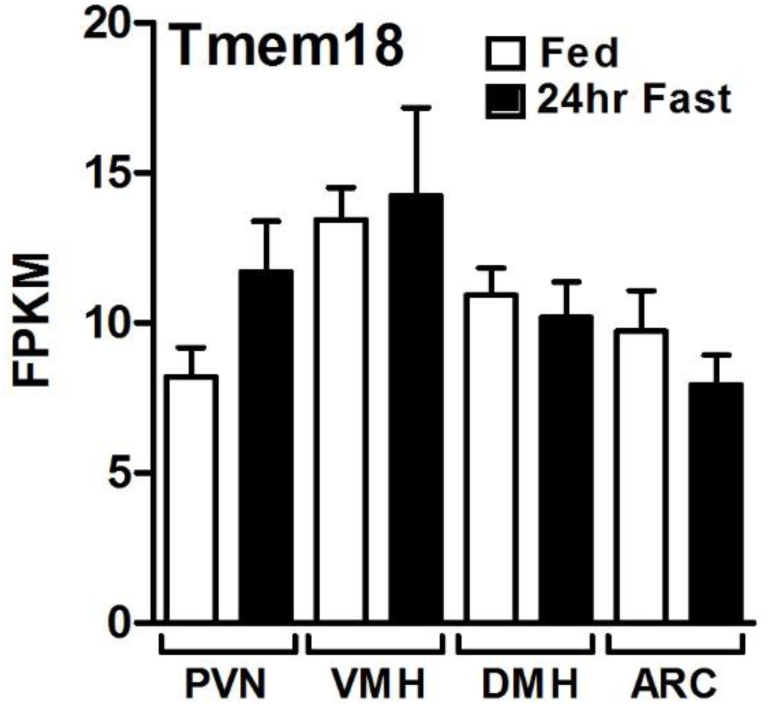
Gene expression of *Tmem18* in laser captured hypothalamic nuclei from *ad libitum* fed and 24 hour fasted wildtype mice. White bars show fed mice, black bars show mice fasted for 24hrs. PVN, paraventricular nucleus; VMH, ventral medial hypothalamus; DMH, dorsal medial hypothalamus; ARC, arcuate nucleus.

**Supplementary Figure 4.**
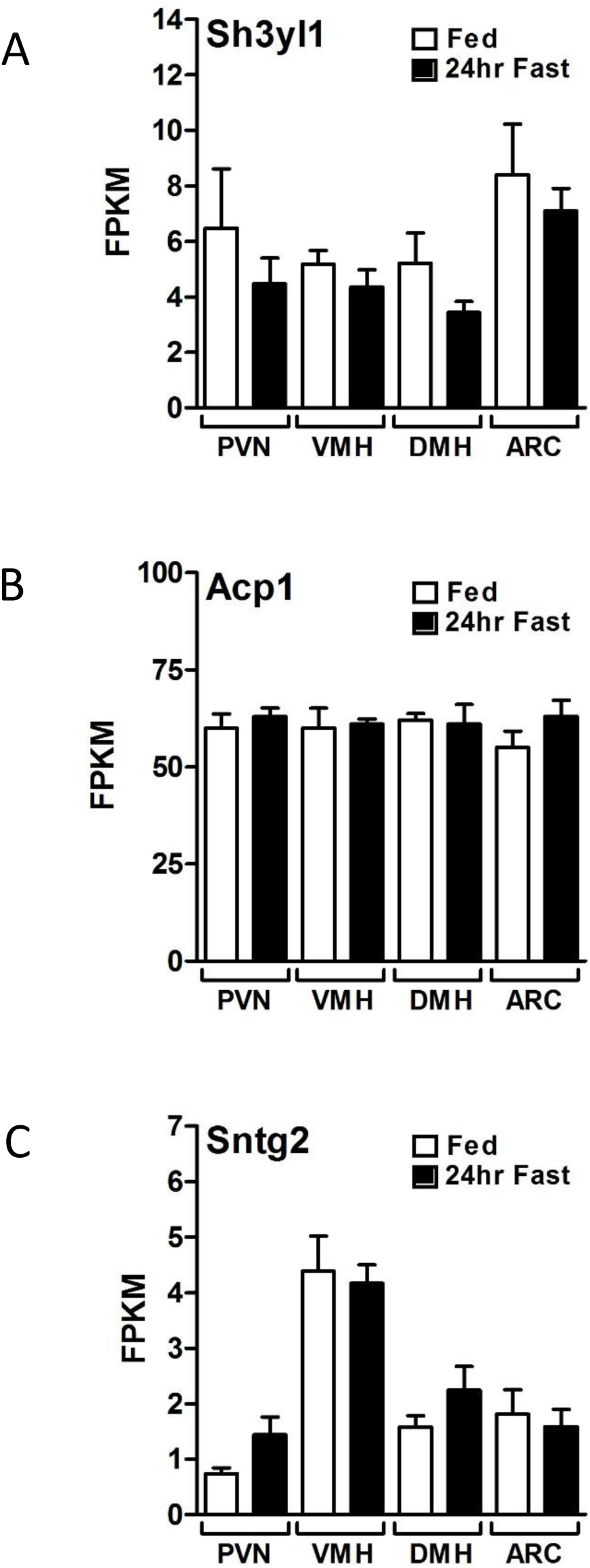
Expression of genes neighbouring *Tmem18* in laser captured hypothalamic nuclei from ad libitum fed and 24 hour fasted wildtype mice. **(A)** *Sh3yl1*, **(B)** *Acp1*, **(C)** *Sntg2*. White bars show fed mice, black bars show mice fasted for 24hrs. PVN, paraventricular nucleus; VMH, ventral medial hypothalamus; DMH, dorsal medial hypothalamus; ARC, arcuate nucleus.

**Supplementary Figure 5.**
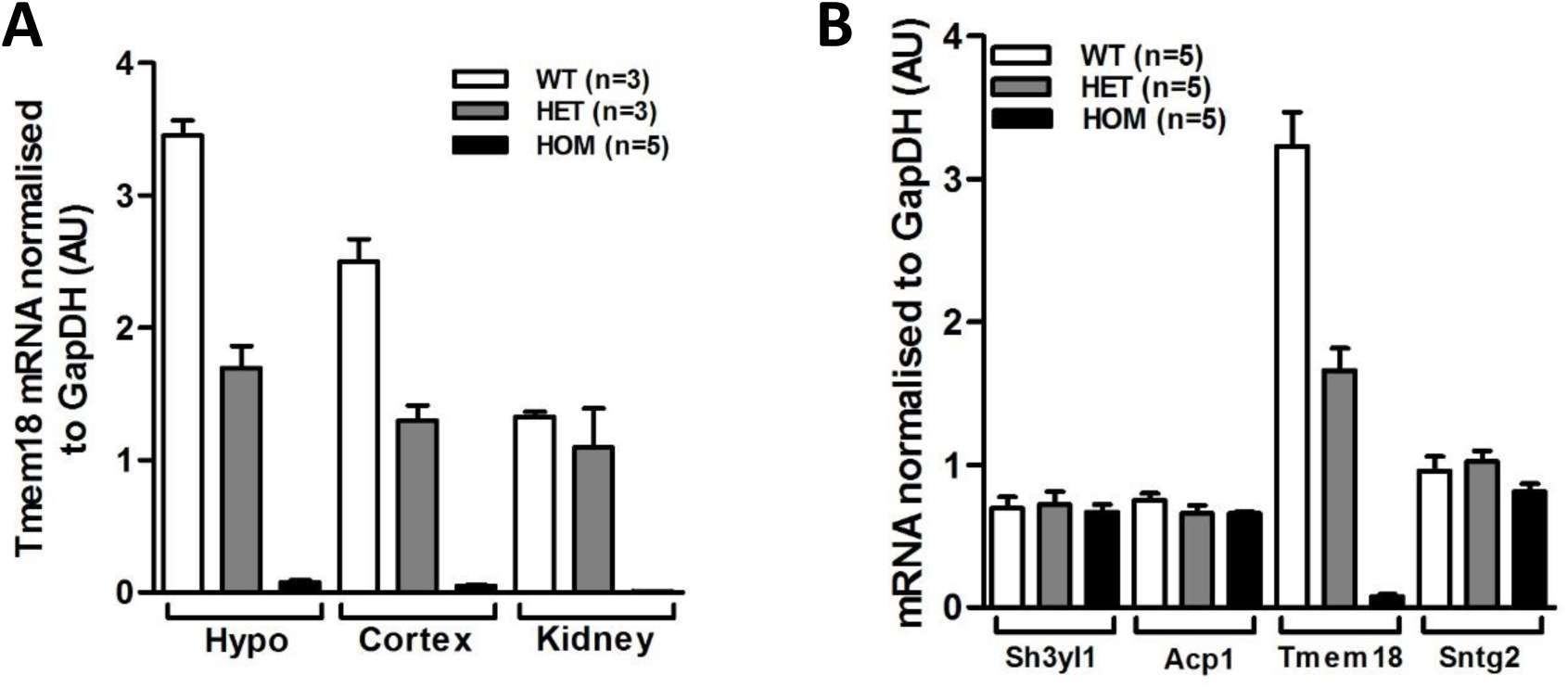
Validation of the *Tmem18tm1a* model. **(A)** Gene expression of *Tmem18* in the hypothalamus, cortex and kidney of WT (white bars), heterozygous (grey bars) and homozygous (black bars) mice, normalised to expression of GAPDH. **(B)** Gene expression of *Tmem18* and its neighbouring genes *Sh3yl1, Acp1* and *Sntg2* in the hypothalamus of WT, heterozygous and homozygous mice, normalised to expression of *Gapdh*. Data are expressed as mean ± SEM.

**Supplementary Figure 6.**
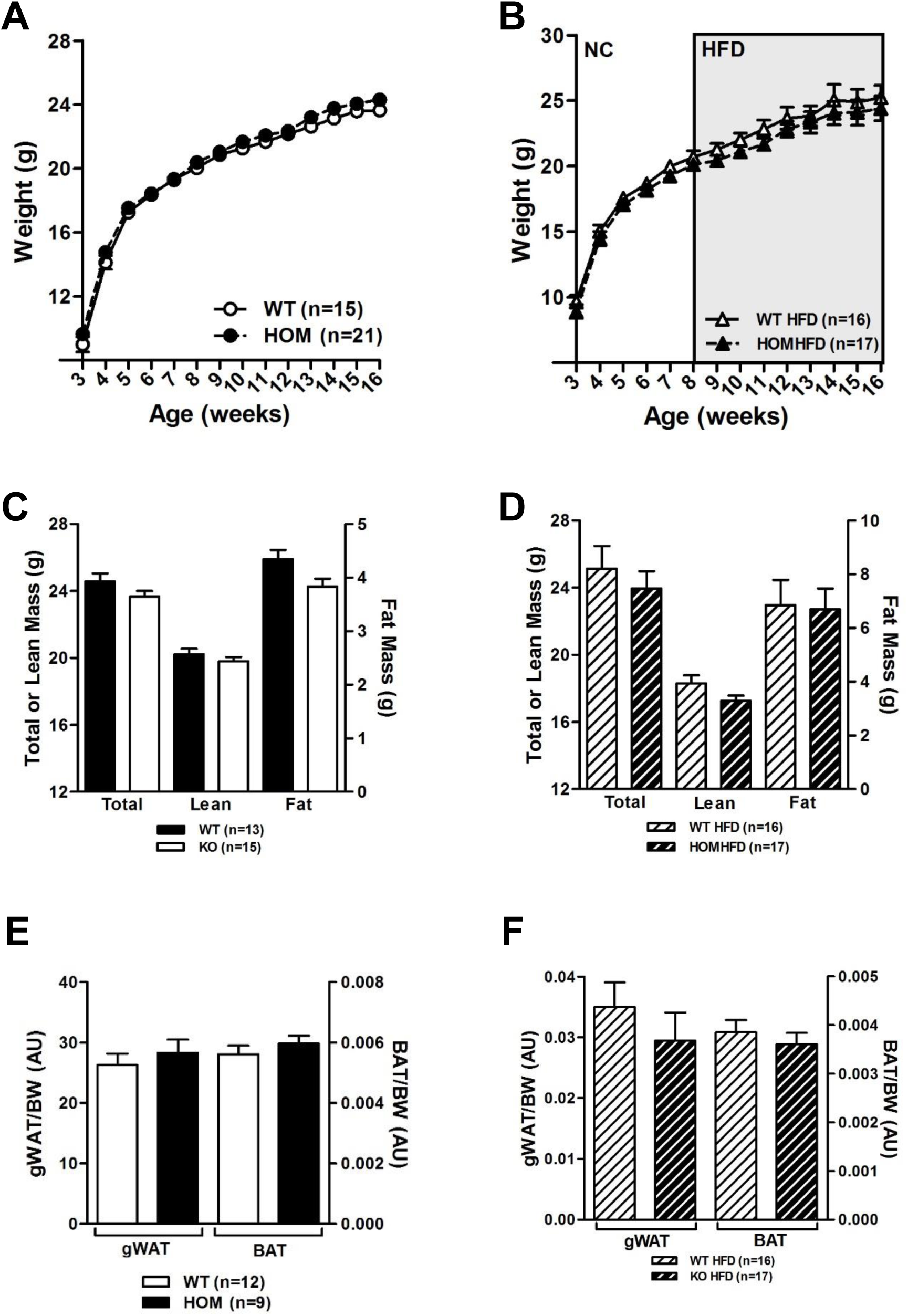
Effect of loss of expression of *Tmem18* on body weight, length and composition in female mice maintained either on a normal chow diet or 45% HFD. **(A)** Body weights of *Tmem18^wt/wt^* (WT) and *Tmem18^tm1a/tm1a^* (HOM) female mice on normal chow. **(B)** Body weights of WT and HOM female mice placed on a high fat diet (HFD) at 8 weeks of age. **(C)** Body composition analyses showing total, lean or fat mass of 14-week old WT and HOM female mice on normal chow. **(D)** Body composition analyses showing total, lean or fat mass of 14-week old WT and HOM female mice placed on HFD at 8 weeks of age. **(E)** Weights of gonadal white adipose tissue (gWAT) and brown adipose tissue (BAT) of WT and homozygous mice at 18 weeks of age, normalised to body weight. **(F)** Weights of gonadal white adipose tissue (gWAT) and brown adipose tissue (BAT) of 18-22 week old WT and homozygous mice fed a HFD from 8 weeks of age normalised to body weight. Data are expressed as mean ± SEM. WT mice are represented by white symbols and bars. HOM mice are represented by black symbols and bars. Time-course data were analysed using the repeated measures ANOVA model. p-values for C-F were calculated using a two-tailed distribution unpaired Student’s t-test and established statistical significance as follows * p<0.05, ** p<0.01, *** p<0.001 vs WT mice.

Supplementary Table 1;List of proteins identified in cells expressing FLAG tagged TMEM18 by immunoprecipitation using FLAG antibody.

